# Structure-property machine learning models with predictive capabilities for glycans in food

**DOI:** 10.1101/2023.11.12.566488

**Authors:** Tarini Naravane, Gabriel Simmons

## Abstract

Structure-property predictive models for food aim to decipher the complex relation between the physical shape of a molecule and its physical properties and/or the functional role of the molecule in a product formulation. Our focus in this paper is the modeling of glycans (i.e., carbohydrates), which are not only abundant in food, but essential to both food production and, more importantly, human health. In our study, we use regression methods to generalize the relationships between the structure of starch (e.g., chain length and composition of protein and amylose) and a range of its properties (e.g., gelatinization temperature, time series viscosity data, gel consistency, and sensory texture) for 301 samples of rice. Our results indicate that the structure-composition data is a significantly better predictor of sensory mouthfeel than the physical properties, even though the latter is typically used in experimental research. This demonstrates the potential to harness several structure-property relationships and we provide strategies to accelerate further experiment-based research.

## 1 Introduction

### 1.1 Need for structure-property research in glycans

Glycans are ubiquitous – commonly known as “carbohydrates”, they are the most abundant class of molecules in nature. They are responsible for various biological activities significant to human health ^1–3^ and, as such, have been a target in food engineering for human nutrition and product formulation. This prominence has motivated research on the relationship between the structural composition of glycans and their physical properties in food formulation. We refer to the structure as defined in the glycobiology text ^4^(and shown in Figure 2), “the primary structure of a glycan is defined by the type and order of monosaccharide residues, by the configuration and position of glycosidic linkages, and by the nature and location of the non-glycan entity to which it is attached.” This definition aligns with recent advances in analytical methods that allow for the detection, identification and characterization of glycans in food^5–7^. For example, in Philips et al.^8^, these measurements have been applied to profile glycan composition at various stages in the ripening of bananas. They correlated the composition at different stages of ripening to various organoleptic and nutritive properties of taste, texture, and dietary fiber. This early work was impactful in revealing the compositional basis for the ripening process but was not aimed at deciphering the relation between glycans and bulk food properties. We hypothesize that leveraging high-resolution glycan composition data will offer performance gains over current practices for a variety of food engineering predictive tasks. This study tests our hypothesis in the setting of rice cooking.

### 1.2 Potential of machine learning models to predict the property from structure

Over the past several years, machine learning (ML) techniques have shown promise in modeling complex relationships between polymer structures and their behaviors. The most prominent example is AlphaFold^9^, which was made possible after two decades of collective effort in building a dataset linking protein sequence to 3D conformation. The breakthrough success of AlphaFold was a stepping stone towards better models of protein function/property, which have since accelerated research in areas such as protein-protein interactions^10^ and drug discovery^11^.

Unlike for proteins, computational techniques for glycans are still nascent, specifically in terms of large datasets and wide applicability. However, early work in (ML) and other computational methods including molecular dynamic simulation (MDS) are being recognized^12,13^. MDS approaches have proven capable of predicting polymer conformation and properties^14–16^, although their utility is limited to short-chain polymers of sizes ranging from 2-25 units due to the computational intensity of the method. Other ML methods have been used to predict properties of immunogenicity and pathogenicity^17^ for single glycans from structural information. Importantly, prior work has not explored the utility of ML in relating molecular-level glycan information to bulk food properties. At present, glycan datasets are considerably smaller than the volume of data that enabled the AlphaFold breakthrough for proteins ^12,18^. While “AlphaFold for glycans” is still far away, early work in glycan predictive modeling encourages us to explore the utility of ML for predicting the properties of glycan-based foods.

### 1.3 Demonstrating the potential of ML models for glycans

Glycans are a major constituent in grains, vegetables, and legumes^19^, and research seeks to understand their roles in food formulation^20^. Generally, a glycan structure is characterized by repeating units composed of a monomer(s), linkages, and optional functional groups^4^.As stated earlier in **Sections 1.1 and 1.2**, the scope of glycans is vast and the analytical progress in discovery is very recent. In contrast, starch is a specific glycan with a long history of analysis and research and continues to be studied due to its diverse physical properties. We therefore selected starch as the representative glycan for our case study. We now give a brief description of the scope and aims of our case study. The dataset is for samples of rice and the data per sample includes the chain length distribution (size exclusion chromatography data) which represents the structural features, content of protein and amylose, the physical properties relevant to the processing quality of rice, and the sensory properties of cooked rice. The architectural overview of the data and models is illustrated in **Figure 1**. The models developed in this study address the following research questions (RQs):

**RQ1:** Is size exclusion chromatography data and content data sufficient to predict both physical and sensory properties of cooked rice?
**RQ2:** Does gelatinization temperature and gel consistency information improve predictive performance over only the chain length and content data (tested in RQ1)?
**RQ3:** What features (structural or otherwise) are the most informative for each prediction?

**Figure 1:**
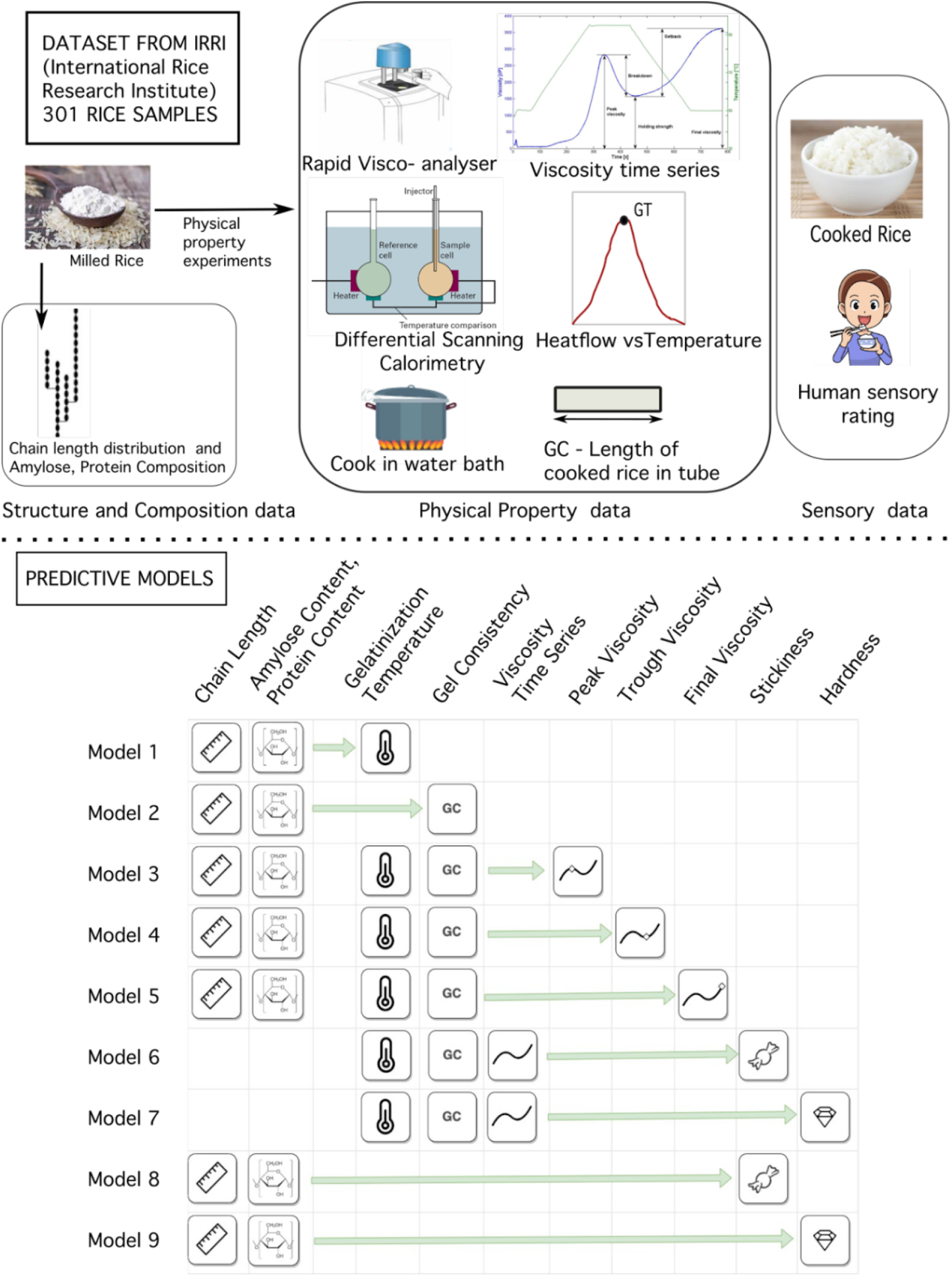
Data and Models. The data is from prior research, ^30^ for 301 samples of rice based on the indicated methods in the upper portion of the figure. 9 Models are trained based on this data as seen in lower half of the figure. Detailed explanation for the data and the models is in section 3.

These research questions address our main thesis that the key to predicting the properties is the detailed structural data. We also dive deeper into the structural features to gain insight into the molecular dynamics responsible for the properties. **Section 2** provides a background of prior research on starch, and its relevance to our methods. The data and methods used in this study are explained in **Section 3**. Our results in **Section 4.1** indicate that structure-composition data is a better predictor of final product texture properties than pasting behavior, contrary to the approach commonly taken in food product formulation^20,21^. Furthermore, the pasting characteristics and other physical properties can themselves be predicted from structure-composition data. The result from feature engineering in **Sections 4.2** and **4.3** identifies the most useful features for each predictive task. Finally, in **Discussion** we suggest strategies to guide future experimental studies towards improving structure property models (**Section 5.2**).

## 2. Background

### 2.1 The Importance of Starch

Food sources of starch like tubers and grains provide up to 30% of daily dietary calories^21^ and starch constitutes 70-90% of the composition in these foods^22^. Starch is structurally simpler than other glycans, consisting solely of a repeating glucose unit with no additional charge. Despite its simplicity, starch is also expressive - differences in its structure across food groups like grains, legumes, and tubers manifest a range of properties essential to food formulations such as binding ability, textural smoothness, and stickiness^23–33^. While our study uses data specific to rice, the features used to train our models are not specific to a single food group (since starch is the shared building block for a host of widely-consumed foods).

### 2.2 Starch Morphology

The structure of starch across units of scale starting from the glucose monomer is illustrated in **Figure 2**. It is composed of two fractions; neutrally-charged chains of glucose, arranged linearly as amylose (AM), and in a branched structure as amylopectin (AP). The consensus among researchers is that the degrees of polymerization (DP) of amylopectin is usually between 9 and 24 up to 100^24,34,35^ while amylose has longer chains. These polymers assemble in an arrangement of crystalline regions composed of branches of amylopectin forming helices due to the strong attractive intermolecular forces, with longer chains forming more stable helices. The amorphous regions are composed of amylose where some long chains of amylose might also form helices. The long-chain helices, whether amylose or amylopectin, are positively correlated with greater crystal stability^24,34,36^. The crystalline and amorphous regions repeat in concentric rings **(Figure 2C)**, with several of these arrangements packed into a granule **(Figure 2D)**. This highly ordered native arrangement is disrupted by food processing operations like milling, heating, or soaking in water.

**Figure 2:**
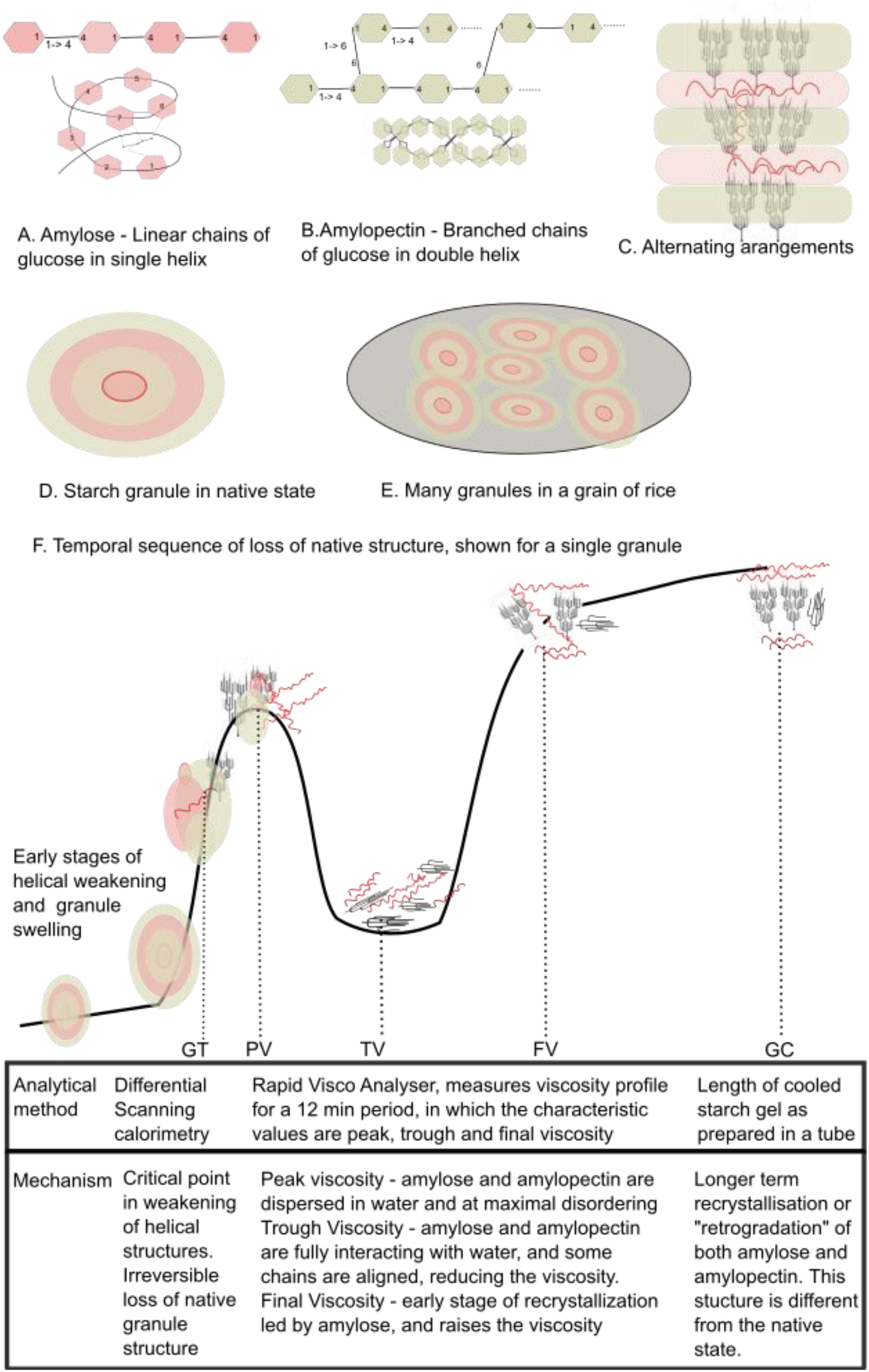
Illustration of the starch morphology in the native state (A-E), and the mechanistic changes to alternate physical arrangements(F). It is important to note that the chain lengths do not change, it is only the state of ordering that changes. The hexagon in A and B represents the glucose monomer. **A**. Amylose is the starch fraction where the monomers are connected by α1-4 glycosidic linkage in a linear chain. Due to the intramolecular forces and the linkages, the linear chain takes on a single helix configuration. **B.** Amylopectin is the starch fraction where the monomers are connected through α1-4 linear and α1-6 branching linkages. The branched chains twist in a double helix. **C.** Amylopectin is in the crystalline region and amylose in the amorphous region.Sometimes the long chains of amylose form a double helix.. The crystalline and amorphous bands alternate and form a granule as seen in **D.** The core is amorphous. **E.**Many granules pack into the grain of rice. **F.** Mechanistic changes leading to loss of crystalline structure and transforming into different arrangements. Note that these are not all measured based on a single sample through time.

### 2.3 Mechanisms driving various physical rearrangements, and properties

Food processing operations, including temperature changes (both heating and cooling) and the addition of water disrupt the native structure of starch matrix The temporal sequence of the various physical arrangements^3^ starting from the native state is illustrated in **Figure 2**. This temporal behavior varies with the composition of the starch matrix. Longer AM or AP chains form more stable helices, requiring greater energy for bond disruption, ultimately leading to higher gelatinization temperatures^24,34,35^. Mechanistically, greater helical entanglement also has the effect of reduced tendency for swelling^36^. When the native starch granule is exposed to water and increasing temperatures, the loss of helical bonding,allows water to enter the granule, leading to further weakening^37^. Upon reaching gelatinization temperature, the bonds have weakened to the extent that the concentric ring structure is no longer present. There are various explanations for the extent of dissociation of the crystalline and amorphous regions that lead to gelatinization; we refer the reader to a recent review by Tetlow and Bertoft^38^. After gelatinization, AM and AP polymers disperse and interact with water. This interaction leads to different values of viscosity depending on the applied temperatures, and this information is important in temperature settings for food processes. The relevant range of viscosities is studied by a standardized analytical test (rapid visco analyser)^25,26,29–31,39^ that measures the viscosity at 4 second intervals for a 12 minute period, where the sample solution is treated to a temperature profile of increasing temperature up to a maximum which is held constant and then decreased. Peak viscosity is the viscous effect at the peak temperature, where amylose and amylopectin are fully dispersed in water and maximally disordered. Higher amylose content typically corresponds to lower peak viscosity, since the amylose lipid complex resists swelling and amylose tends to form linear chains which slide over each other in solution^40^. When the temperature is held constant, the viscosity falls as amylose and amylopectin chains are aligned while still interacting with water, and this is the trough viscosity. The starch polymers are fully dispersed at this point^41^. Then the temperature is reduced leading to an early stage of recrystallization – or “retrogradation”, and the resulting crystalline pattern is different from that of the native crystal structure^36^. This starts with the shorter chains of amylose^42^ in the early stages and followed by amylopectin. This mechanism is observed as an increase in the viscosity up to the final viscosity, which is proportional to the amylose content. Long chains of amylose take longer to retrograde, and are associated with greater hardness of the cooked rice^24,36^.

Stickiness is inversely related to amylose content and positively related to amylopectin that have a delayed retrogradation^34,43,44^. Retrogradation effects are studied for their detrimental effect on sensory properties - for example the hardening of baked goods^45^.

However, there are gaps and limitations in prior research in discovering the relationships of structure to the properties, as examined in recent review papers. Hamaker explains that the cumulative structural knowledge of starch lacks details of the internal architecture of amylopectin which limits structure-property applications in food formulation^46^. Tetlow and Bertoft ^38^ provide an elaborate description of the starch granule in sections 2-6 of their paper as a basis for their critique and proposed solution. They point out the inadequacy of frequently reported features in prior research such as short, medium and long branches of amylopectin, and amylose content by citing contradicting observations or unexplained differences in phenotypes. The authors then reference their prior research^47^ where Bertoft proposed the backbone branching structure of amylopectin (also called interblock chain lengths) as an important morphological feature which addressed their critique. The inadequacy of morphological features was also investigated by Tao et.al^48^, and they reported the influence of the molecular size of amylopectin on the viscosity properties.

In our opinion, another gap in research is relating high resolution structural features to both the granule morphology and the properties. This would not only connect all the concepts across scales of resolution, but also provide greater precision to structure-property models. The latter objective has also been suggested by Yu et. al ^49^. We explore solutions for this gap in **Discussion Section 5.2**.

### 2.4 Relation of domain knowledge to experimental hypotheses

The main features of the starch morphology described in the previous sections are the chain lengths, the branching structure at the nanometer scale, and crystallinity at the scale of the granule morphology. These are reflected in the dataset (details in **Section 3.1**) as; (i) the chain length features are provided by the SEC data, (ii) the amorphous fraction of the crystal addressed by the amylose content, and (iii) the branching structure and crystallinity are represented by observational data of gel consistency and gelatinization temperature. Our central hypothesis is based on the fact that the structural components at the nanometer scale remain constant during processing but acquire different physical arrangements each connected with a specific property. However, since the dataset does not contain branching structure information at the nanometer scale, we use the data assumptions mentioned earlier in this section and address the hypothesis in two parts (**RQ1 and 2 in Section 1.3**). First, we test whether the chain length distribution and content data is a versatile predictor. Then to test whether data on branching and crystallinity aspects improves prediction, we use the empirical measurements of gel consistency and gelatinization temperature. We compare the predictive performance based on chain length and content data alone and with the addition of gel consistency and gelatinization temperature. Further details of model design are in the **Methods Section 3.2 and 3.3**.

### 2.5 Case study and related work

The authors Buenafe et. al. from the International Rice Research Institute, show the promise of predictive methods for understanding how starch structural data and physical and sensory properties are related^39^. The food samples in their dataset are varieties of rice. The data per sample includes the chain length distribution (measured by size exclusion chromatography) of starch, the physical properties relevant to the cooking of rice, and the sensory properties of cooked rice. Our case study uses the same data, and **Section 4.1** has the detailed description. The approach we take builds on this prior work in several important ways. First, the method proposed by Beunafe et. al. predicts the cluster index for an unseen sample to indicate similarity to the previously identified clusters of rice samples but does not directly predict physical or sensory properties of the unseen sample. We address this limitation by reframing the predictive task to predict the values of the physical and sensory properties directly.

Another limitation is that Buenafe et. al. do not use the high-resolution chain length distribution data for the clustering method, and instead use a reduced version by summing into 5 groups. We speculate that this is due to the challenge of high dimensionality presented by the 500 SEC features. We include this complete data in our predictive models and show several methods that can be used to identify and overcome the dimensionality issues. In addition, we go a step further with our modeling approach that allows us to interpret feature importance values in relation to prior knowledge about the mechanistic behavior of starch polymers.

## 3. Methods

### 3.1 Data

#### Data samples and features

We used the dataset from the study (referenced in **Section 2.5**) by Beunafe et al. at the International Rice Research Institute (IRRI)^39^ which includes three types of data: composition, physical, and sensory, as shown in **Figure 1**. The composition data includes amylose content (AC) and protein content (PC). The structural data was measured by size exclusion chromatography (SEC) and reported for 500 degrees of polymerization (DP) values in the range from 5 to 12,000. The physical properties consist of gelatinization temperature (GT) measured by DSC, gel consistency (GC) measured as the length of the starch gel prepared in a tube after heating (followed by a hour of cooling), and time series viscosity data using a rapid visco analyser (measured every four seconds for a 12-minute period). The sensory data included 13 mouthfeel descriptors and was collected for 100 samples of rice. For this case study, we focus on hardness and stickiness, since these are the most widely studied sensory characteristics across literature and fundamental to other properties like cohesiveness and toothsomeness^43,44^.

#### Data selection and qualitative observations

The original study data contains 301 samples of rice, of which 100 had sensory data. We discarded all samples with missing data, resulting in a dataset of 231 samples for the models predicting viscosity, gelatinization temperature, and gel consistency. Of these, only 72 samples had sensory data and were used for the models predicting sensory characteristics. The data visualization is shown in **Figure 3**. Viscosity time series data shows that the samples vary in the gradient and overall profile. There are peak viscosities at roughly two distinct times; the first is just after 200 seconds with a peak value of 2500 centipoise and the second is between 300-400 seconds with a peak value ranging from 2000-200 centipoise. The trough viscosity has a wide range, though it all occurs at 500 s, after which it increases. Similarly, there is also a wide range in final viscosities, and it appears that the viscosity gradient from the trough viscosity varies greatly. This suggests that some samples have a faster rate of retrogradation. The histogram of the amylose content (AC) histogram shows that there are a few samples with almost no amylose, otherwise the AC variation is small for the majority of samples. The histogram of protein content (PC) has a wide range, and usually for starchy grains, protein content above 10% is considered high. Gel consistency and gelatinization temperature have a wide range of distribution which does not have a uniform shape. Hardness and stickiness also have a wide range implying the variation in rice samples perceived by consumers. The subset used in this study are provided in the Supplementary Information File, and we refer the reader to the cited publication by Beunafe et al. ^39^ for the complete dataset.

**Figure 3:**
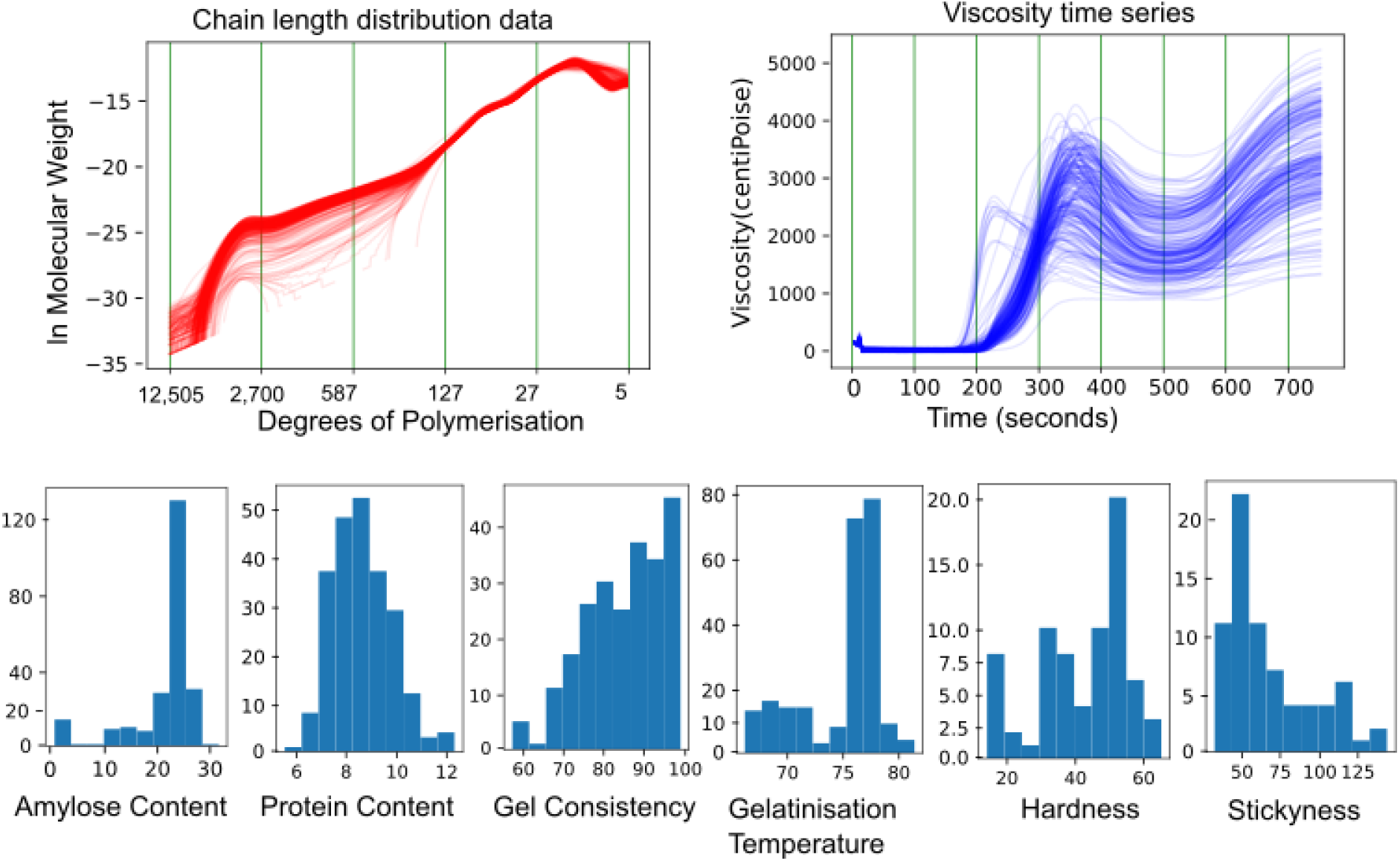
Illustrating the data for the 231 samples of rice. There are two types of data. The distribution is the chain length distribution data and viscosity data which is shown by line profiles per sample. The rest of the data is discrete, i.e. a single value per sample and shown by histograms.

### 3.2 Predictive Models

We designed a total of nine models, as shown in **Figure 1**, to address the research questions in **Section 1.3**. Models 1 and 2 predict gelatinization temperature and gel consistency from the composition and structure data. Models 3-5 predict peak, trough, and final viscosities from the composition and structure data, with some variants also using gelatinization temperature and gel consistency.

Models 6 and 7 predict sensory properties of stickiness and hardness from the composition and structure data. Models 8 and 9 predict stickiness and hardness from the physical properties – viscosity time series, gelatinization temperature, and gel consistency. Results from Models 8 and 9 are compared to Models 6 and 7 to test whether structure and composition data is a better predictor of sensory properties.

### 3.3 Feature engineering and feature set selection

We used two different strategies in setting up the predictive feature set for a given prediction target. The first was to address a likely problem of high dimensionality of the distribution-type data, as identified in **Section 3**. This applies to the chain length data with 500 DP features, and the viscosity time series data with 188 measurements over a 12 min period. We compared two techniques for dimensionality reduction feature aggregation for the chain length data, and interval sampling for the viscosity data. Ten variations of chain length data were generated – one at full resolution and an additional nine variants aggregated at varying bin sizes. Twenty-five variants of the viscosity data were generated: as is, and also as a time derivative to smooth out the curve for four time intervals (3, 5, 7, 10 seconds), combined with sampling at five different sampling rates - i.e every nth value (1, 3, 6, 8, 10). The viscosity time series data for the time interval variants is the derivative value instead of the original measured viscosity value, and differs for each of the four time intervals. At a certain interval, the value (at each of the 188 points) is the gradient(dv/dt) for that interval.

For each predictive target we tested various predictive feature sets, and the variants of the chain length data and viscosity data were a part of these feature sets. The feature sets with physical property features include GC and GT in four different variants: both GC and GT, either GC or GT, or neither. The feature sets with composition features include AC (amylose content) and PC (protein content) in four different variants: both AC and PC, either AC or PC, or none. For the viscosity prediction models (Models 3-5) where we use both physical properties and compositional features, we use only these five combinations: AC, PC, GC, GT, or AC, PC, GC or AC, PC, GT or AC, PC, or none. This is designed with the intention to test for the inclusion of GC and GT as predictive features as explained in **Section 1.3**. Putting all this together, the feature set variants for the nine predictive models are listed below, and illustrated in **Figure 1**.

1. Gel consistency (model 1) gel temperature (model 2), stickiness (model 6) and hardness (model 7). Total of 40 variants. Ten variants are for the chain length data by bins of size 1, 5, 10, 15, 20, 25, 30, 35, 40, and 50, and four variants for the content data: AC and PC, only AC, only PC, and neither.
2. Peak, trough, and final viscosity (models 3, 4, 5). Total of 50 variants. Ten variants are for the chain length data by bins of size 1, 5, 10, 15, 20, 25, 30, 35, 40, and 50. 5 variants for the other features are AC, PC,GC and GT;AC, PC and GC;AC,PC and GT;AC and PC, and none.
3. Stickiness (model 8) and hardness (model 9): 100 variants. Twenty-five variants are for the viscosity time series, four variants for the content data: AC and PC, only AC, only PC, and neither.

### 3.4 Model Training

The approach to model building had two main steps: hyperparameter tuning and feature selection.

#### Hyperparameter Tuning

Random Forest models were trained for all the variants in each predictive task using the Random Forest regressor from Scikit^50^. We performed hyperparameter tuning to identify an appropriate hyperparameter configuration for each model, through a grid search cross validation method for a range of hyperparameters (range specified in the **Supplementary File)**. Hyperparameter configurations were selected on the basis of mean absolute error (MAE). The grid search results for all variants are provided in the **Supplementary File**. The MAE values are converted to a mean absolute error percentage (MAPE) by dividing the MAE by the average value. MAPE is scale independent and is suitable for comparison of the predictive performances.

#### Model Selection

A subset of all trained models (with hyperparameter configuration) per prediction target were selected for the next feature selection method, since it is computationally intensive. We explain the selection criteria through the example of the peak viscosity prediction; the results of the 50 trained models (**details in Section 4.3**) were grouped by the five discrete-feature combinations (AC, PC, GC and GT; AC, PC and GC; AC, PC and GT; AC and PC, and none). In each group the model configuration^4^ with the lowest MAE value was selected. We also performed feature selection for model configurations using the SEC data with bin size 5. These models did not have the lowest MAE; we include them to compare model performance for high- and low-resolution data (see Section 4.2).

#### Feature Selection

We used the Sequential Feature Selection method (SFS) in backwards mode, from the ML Extend library^51^. The feature selection method associates every feature (of the predictive feature set) with its importance in lowering the prediction error.

Specifically, this method starts by considering models trained on all *n* features, then all possible combinations of *n-1* features, and so on until only a single feature is considered. The importance of each feature is determined by its rank. Features removed at the nth step are considered rank 1, features removed at the (n-1)-th step are considered rank 2, and so on.. **Figure 5** shows results from this method – removing features incrementally improves the performance until a peak, after which removing additional features degrades performance. It is important to note that the feature rank does not indicate a positive or negative correlation between the feature and the prediction target. We run ten replicates of this method to address any randomness associated with the results and focus on the consistent features across the replicates when we report the results (see **Figure 6**).

#### Best Model Configuration

The best model configuration for each predictive task is the model configuration resulting in the lowest MAE for that predictive task. The performance results for the best model configuration for each predictive task are presented in **Table 1**. The optimal sets of features for each task are presented in **Table 2**. Optimal hyperparameter configurations are in the **Supplementary File**. The naive ML baseline for comparison was the average value of the ground truth values for each target variable.

## 4. Results

### 4.1 Structure and composition data are versatile and outperform physical features in predicting sensory attributes

The performance metrics for the best models 1-9 are in **Table 1** (see **Section 3.4** for best model selection). These metrics present two key inferences; first, the structure and composition features are better predictors of sensory properties relative to physical property features, and second, the physical properties can themselves be predicted from structure and composition. In combination, these results indicate that structure and composition features are versatile in their predictive capability. Furthermore, all predictive models have an average error 42.43% lower than the baseline models.

Predictions of sensory properties (models 6-9 for stickiness and hardness) from structure and composition features resulted in 27% lower MAPE in comparison to prediction based on physical property features. More specifically, MAPE for prediction from structure and composition is 15.84% for stickiness and 11.31% for hardness. The MAPE for prediction from physical property features is 20.20% for stickiness and 16.62% for hardness. Both approaches outperform the naive baseline, which achieves 35.09% MAPE for stickiness and 25.95% for hardness. Physical properties can themselves be predicted from structure and composition data. Complete details of the performance metrics for the predictive and baseline models 1-5 are in **Table 1A**, and here we summarize the relative MAE improvements of the prediction models compared to the baseline models; 10.73% for gel consistency, 40.84% for peak viscosity, 39.89% for trough viscosity, and 45.34% for final viscosity. Relative prediction error for gelatinization temperature was better, but only by a small margin in absolute terms; the prediction model MAE was 1.3, but the baseline model error was already good at 2.93 against an average value of 75.26. Complete coverage of all tested variants is in the Supplementary File.

**Table 1A:** Comparing results from ML model (predicted from the structure and composition data) and the baseline model (which computes the average value). MAE is mean absolute error, and MAPE is mean absolute percent error.

| Model number, predictive target | ML model |  | Baseline (Average) |  | Error Reduction (ML vs Baseline) |  |
| --- | --- | --- | --- | --- | --- | --- |
|  | MAE | MAPE (%) | MAE | MAPE (%) | MAE | MAPE |
| 1. GT | 1.30 | 1.73 | 2.93 | 3.9 | 55.63 | 55.64 |
| 2. GC | 7.70 | 9.27 | 8.63 | 10.38 | 10.73 | 10.69 |
| 3. PV | 297.88 | 10.54 | 503.49 | 18.04 | 40.84 | 41.57 |
| 4. TV | 249.78 | 13.93 | 409.36 | 22.82 | 38.98 | 38.96 |
| 5. FV | 338.92 | 10.25 | 620.03 | 18.74 | 45.34 | 45.30 |
| 6. SBG | 10.77 | 15.84 | 23.86 | 35.09 | 54.86 | 54.86 |
| 7. HRD | 4.87 | 11.31 | 11.16 | 25.95 | 56.36 | 56.42 |

**Table 1B.**
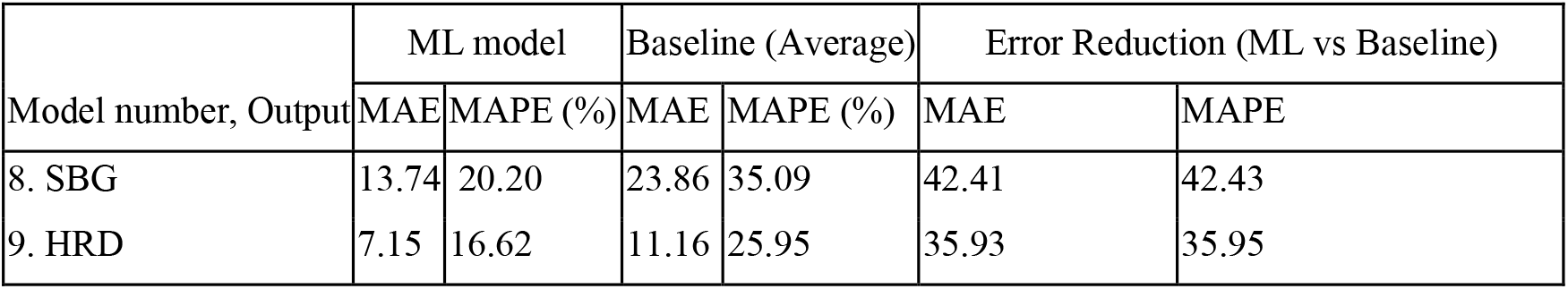
Comparing results from ML model (predicted from viscosity time series, gel consistency and gelatinization temperature) and the baseline model (which computes the average value). MAE is mean absolute error, and MAPE is mean absolute percent error.

### 4.2 High-resolution models outperform low-resolution models in predicting physical properties

In identifying the likely challenges to our approach, we had considered that the high dimensionality of the structural data – which spans more than 500 DPs– combined with low sample size may impede predictive performance. We explored and compared two solutions for dimensionality reduction: low-resolution features via binning and selecting high-resolution features using an iterative feature selection procedure.

For the low-resolution feature approach, we used a binning method to reduce the size of the chain length data (**see Section 3.3**). Then, we trained several models with these different versions of the chain length features to identify the bin size resulting in the lowest predictive error. **Figure 4** shows a lower sensitivity to bin size for viscosity predictions, and greater sensitivity for gelatinization temperature, gel consistency, and sensory properties prediction. Bin size 5 consistently results in lower average error than the full resolution, for all prediction targets except gelatinization temperature. Another consistent trend is that the error decreases for lower resolutions, with the least error in bin sizes from 20 - 30. The error then increases for the lowest resolution at bin size 50.

**Figure 4:**
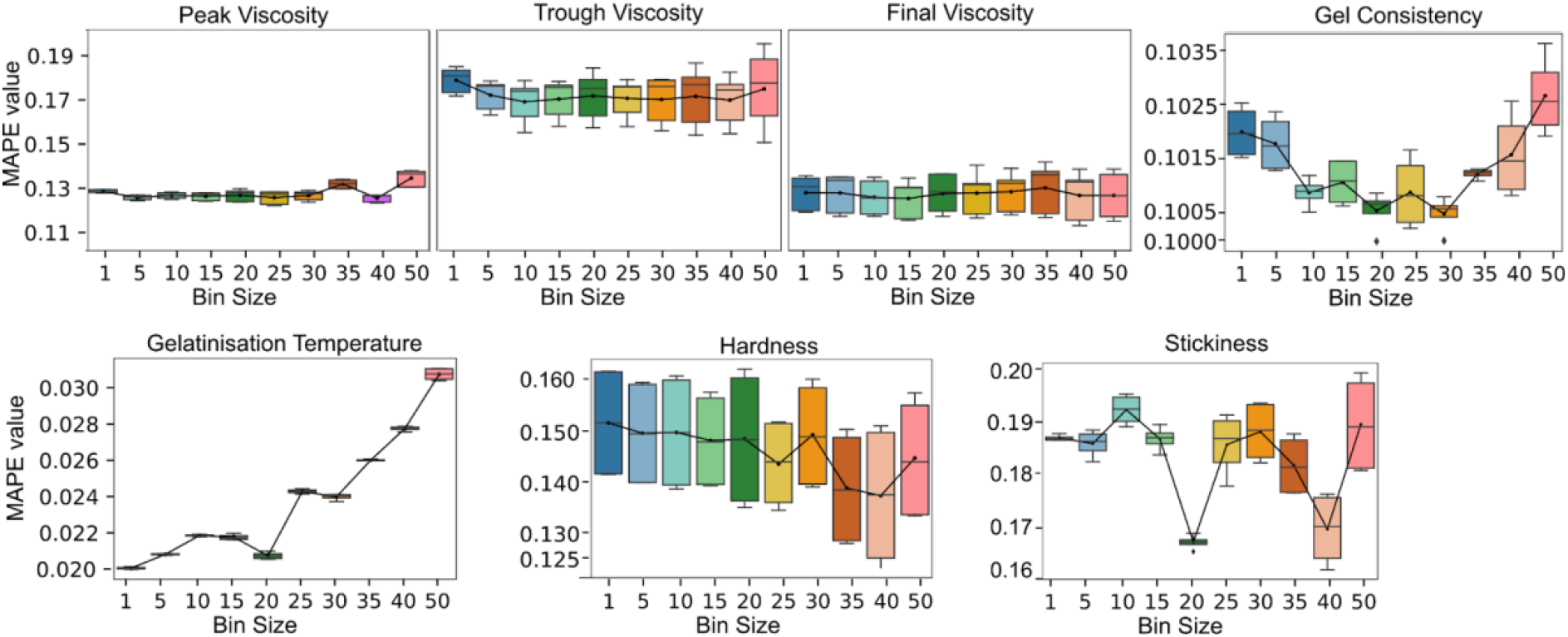
Predictive performance (MAPE) after running hyperparameter tuning step, of model training. Box plot illustrated the results grouped by the bin size of the chain length distribution data. Bin sizes are 1, 5, 10, 15, 20, 25, 30, 35, 40, 50. Full resolution is 500 DP values. Each box plot represents the performance of all the models trained on different discrete features for the same feature aggregation (bin size for SEC). Within each box plot, the performance range is therefore due to the different discrete features sets. The averages are connected to illustrate the trend which is described in result 4.2.

The high-resolution approach is based on the feature selection method (**see Section 3.4**). We run this for both the models based on the high-resolution features at a bin size of 5, and the model based on the bin size identified in the low-resolution approach^5^. We then compare the performances for the full feature set and with the optimal feature set for both resolutions in **Figure 5**. Feature selection improves performance for both the high and low-resolution feature sets Gains from feature selection are greater for high-resolution features for the physical property predictive models (peak viscosity, trough viscosity, final viscosity, gelatinization temperature and gel consistency). The specific features in the optimal feature set for the high-resolution version are listed in **Table 2**.

**Figure 5:**
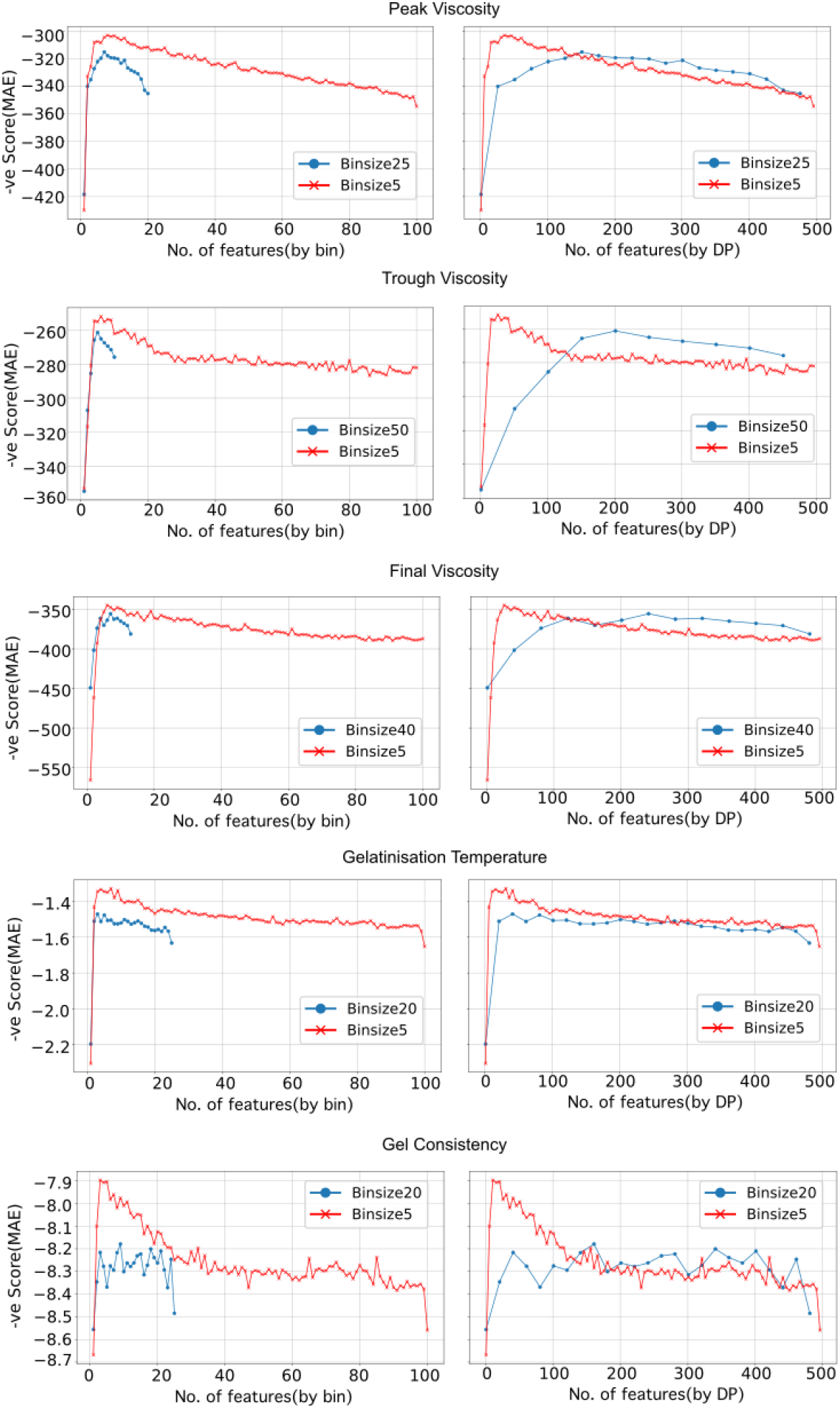
Predictive performance (mean absolute error) from the feature selection method. Overall these figures show gain in performance from the optimal features and is described in result 4.2. Each plot shows and compares the performance for two different resolutions of the chain length data features. The MAE numbers are reversed for convenience of visualization such that the high peak in the curve is the best performance. The legend indicates the bin size used for aggregating the chain length data. The same result is shown through two perspectives on the left and right. The y axis in both views is MAE, and the perspectives differ in x axis only. Essentially the view on the right is a “stretched out” version. **Left :** The X axis is the binned feature. The high resolution models have more features, than the low resolution model. Note that this plot does not have any information about the specific degrees of polymerization for each feature. Also, the aggregated feature b1 for the high resolution is not the same as the aggregated feature b1 for the low resolution. **Right:** The X axis is the DP coverage. The low-resolution plot has fewer bins, and every bin has a much greater feature coverage. Each plot point is positioned at the start of the bin, and the last point covers the last bin even though the trend line ends short of the DP 500 on the x-axis. Note that the predictive features do include the other features (AC, PC, GC, GT), but they are not shown in this plot. For example, if feature 1 was a binned feature, feature 2 was AC, and feature 3 was a binned feature, we only show feature 1 and feature 2. So, although certain features are omitted, the MAE values are preserved by feature rank.

**Table 2.** The predictive features selected across all 10 replicates are listed per property.

| Model number, predictive target | Predictive Features |  |  |
| --- | --- | --- | --- |
|  | Amylopectin Dps | Amylose Dps | Other features |
| Gelatinization Temperature | 7,12, 13, 19, 20, 81-86 | 6300-6700 |  |
| Peak Viscosity | 47-50, 64-80 | 175-200, 300-320, 5000-6700 | GC |
| Trough Viscosity | 32-43 | 162-200 | AC,GC,GT |
| Final Viscosity | 35-44, 70-74 | 167 -200, 5100-5800 | AC,GC,GT |
| Gel Consistency | 28-35, 48-54 | 150 - 186 |  |

### 4.2 Predictive features occupy concentrated regions of the DP space across resolutions

Although we identified that the high resolution features have better performance in result 4.2, we explored the feature ranks at each resolution to understand the reason for the different performances by resolution. The ranks of the optimum features are presented as a heatmap in **Figure 6**. The rank information is given for 10 replicates of the feature selection method for each model at a specific resolution. We indicate the general alignment of features across the resolutions by vertical lines. These aligned features also match in their ranks as indicated by the color codes. As we further analyze this alignment, it is essential to note that each low-resolution feature aggregates more DPs than the high-resolution features. We observe that for any low-resolution feature only a few of the corresponding high-resolution features have a positive rank. This result suggests that the predictive capability of an aggregated feature is influenced by its constituents - what feature ranks does it aggregate. This observation means that not all the aggregated features are powerful simply for being large. The aggregated features that combine one or more predictive high-resolution features are themselves predictive. Based on these observations, although there is a general consistency in the optimum features across resolutions, the low-resolution features are weaker in their predictive capability compared to the high-resolution features.

**Figure 6:**
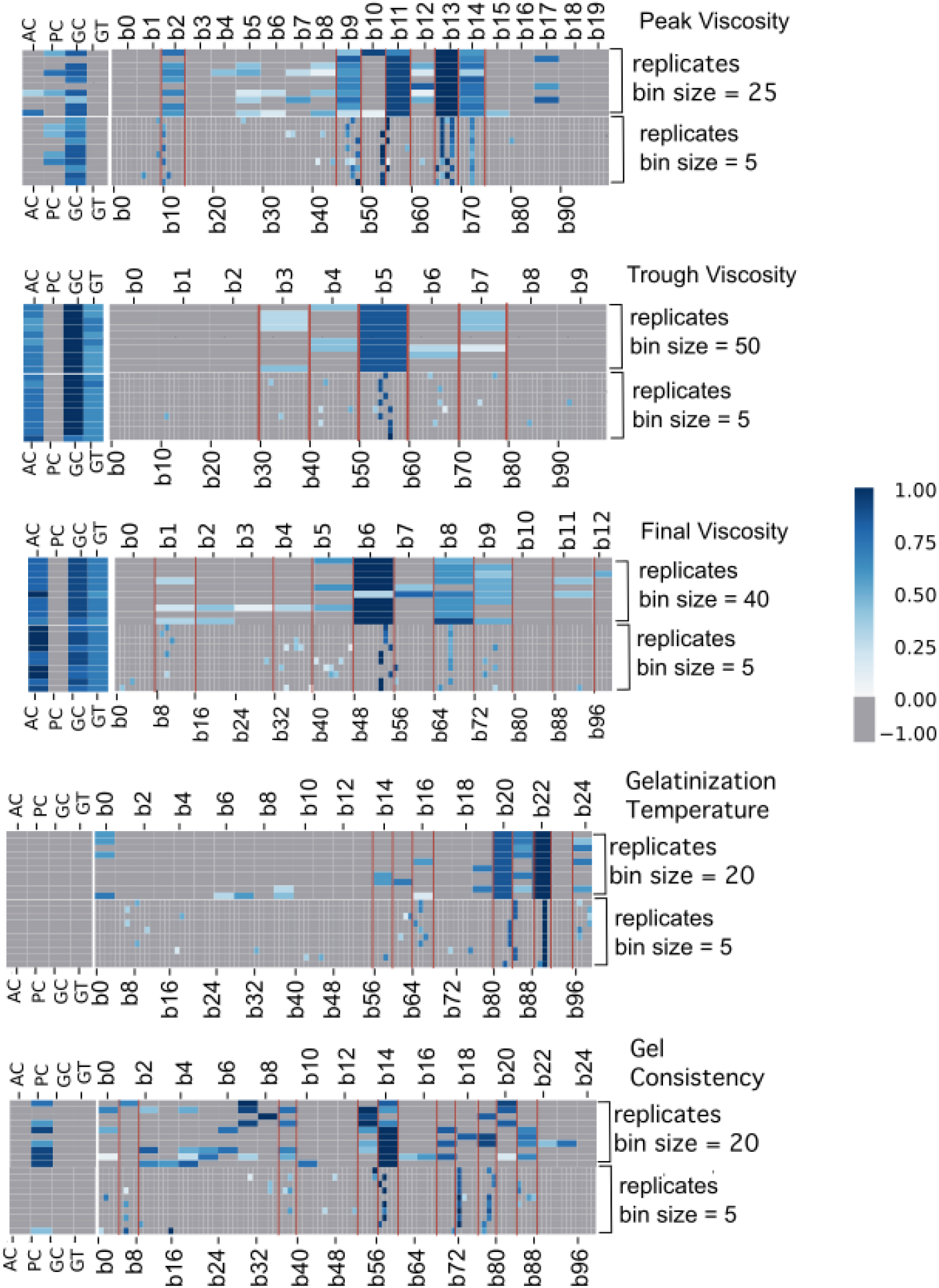
Feature rank results for physical property prediction models. The heat maps compare optimal feature sets for different resolutions of the chain length data, per predicted target. There are 10 replicates per model. For each set of replicates, the bin sizes are indicated to the right. The features by bin number are marked corresponding to the map grid. The actual DP ranges corresponding to the high-resolution bin features are marked at the bottom of the grid. The optimum features are assigned a blue color-code by rank, and all negative ranking features are assigned a uniform gray color. The vertical lines running through the grid indicates the alignment between features across the resolutions.

### 4.3 Crystalline structure information improves predictions of peak, trough and final viscosity

We verify the hypothesis in **Section 2.4**, that additional features on the crystallinity aspects of structure would improve prediction predictive performance. As we are limited by our dataset, we use gel consistency and gelatinization temperature as a proxy for the information on the branching and degree of crystallinity. The predictive performance with these additional features improves the peak viscosity prediction by 2.7%, trough viscosity prediction by 16.83%, and the final viscosity by 10.82%, as shown in **Table 3**. We discuss this topic further in **Section 5.2**.

**Table 3.** Model performance (mean absolute error) for models trained on predictive feature sets that exclude or include gel consistency and gelatinization temperature. Model performance is better (lower error) for models trained with GC and GT.

| Predictive target | MAE (excl. GC,GT) | MAE (incl. GC,GT) | Relative improvement (incl. GC,GT) |
| --- | --- | --- | --- |
| Peak Viscosity | 305.25 | 297.01 | 2.70% |
| Trough Viscosity | 300.35 | 249.78 | 16.84% |
| Final Viscosity | 380.06 | 338.92 | 10.82% |

## 5. Discussion

The results successfully demonstrate that structure and composition data is a versatile predictor of a range of physical and sensory properties. Further, to emphasize the potential of structure-property models, we find that structure and composition data is a better predictor (27% lower error) of sensory mouthfeel than physical property data. This superior performance result challenges the prevalent approach in experimental research that leverages the measurement of pasting dynamics to predict textures properties^29–3152^. We also found that models trained on selected high resolution chain length features led to superior predictive performance and we identified the most predictive chain lengths for each physical property.

### 5.1 Relating chain length features to the mechanistic changes and material properties

Prior research (addressed in **Section 2.3)** has studied details of starch morphology and relationships to the mechanisms and properties with the common objective of making starch behavior predictable, but there are gaps in the knowledge to date [all citations from section 2.3]. Our suggested solution to this gap is to relate high resolution structural features to the morphology and properties. We exemplify our solution by relating the feature importances (**Section 3.3**) to the prior research findings for the gelatinization temperature. The optimal predictive features for gelatinization temperature are amylopectin DPs 7, 12, 13, 19, 20, 81-86, with higher ranks and amylose DPs 6300 --6700 has the lowest rank. Gelatinization temperature is associated with the loss of crystallinity and granule structure, and results from exposure of the granule to heat and moisture for a period of time. The mechanisms responsible are the simultaneous disruption of helical bonds from the heat, penetration of water into the center of the granule and leaching out of amylose. Tao et. al^48^ have shown the correlation of amylopectin DP 6-60 and short amylose branches (DP 270-354) ^6^ to the gelatinization temperature. The amylopectin features agree with the research by Tikapunya et. al which shows the correlation of DP 6-33 to the gelatinization temperature^53^. However, the findings by Li et al. ^42^ on amylose DPs contradicts with Tao et.al, since it shows leaching of long amylose chains though it was unclear whether this observation was at the point of gelatinization or later. In relating these results to our feature results, we find strong evidence for amylopectin from two sources but contradictory evidence for amylose DPs. In general, it is difficult to interpret our results considering current mechanistic knowledge which lacks agreement on relationships between DP features and properties.

### 5.2 Insights to guide future experiments in data generation and modeling

The prior topic in discussion addressed that the branching structure influences the mechanisms and the expressed property, and that this morphological feature has been under researched comparatively. The need for data on the degree of branching and crystallinity as predictive features was also hypothesized (**Section 2.4) and** raised as a research question (**RQ2 in Section 1.3)**. The results in **Section 4.4** confirmed that inclusion of gel consistency and gelatinization temperature, as a proxy for analytical data, improved predictive performance. At the same time, gel consistency and gelatinization temperature hough predictable from the structure and composition data, do not have 100% accuracy. This hypothesis can be more precisely validated by including glycosidic linkage data on the abundances of α1-4 (linear connections) and α1-6 (branching points) bonds ^54^.

## 6. Conclusion

The results from our case study on structure-functional predictive capability are unprecedented, despite the enormous prior existing research on starch including many journals specific to starch. This case study addresses the questions that are studied extensively in research literature; 1. Is there a relationship between the starch structure/composition and the physical and sensory properties and 2. Is RVA and other empirical data indicative of grain quality and/or product attributes. These questions are significant to both breeding crops for starch structure and food formulating for desirable traits in the products. However, there exist no standard models and only a few prior experimental studies have only established statistical correlations or simple linear equations, and on experimental datasets of a few samples^55–57^.

In response to the first question, the physical properties of peak, trough and final viscosity, gel consistency, gelatinization temperature and sensory properties of hardness and stickiness of rice are predicted from high resolution data on the composition and structure. To address the second question stated above, hardness and stickiness are also predicted from the physical properties which had a 27% lower predictive accuracy. Further, we obtain the specific chain lengths of starch polymers that are most predictive of a physical property and compare these with prior domain knowledge assembled over decades of observational studies and discover that there is a lack of agreement on relationships in literature.

Ultimately this study shows that the ability of machine learning methods to learn complex multivariate relationships is certainly applicable in the complex domain of food science and food formulations. Such data-driven analysis can reveal insights that are nearly impossible to discover in hypothesis-driven experiments. This manuscript is a call to creation of standardized datasets of biopolymers in food, for enabling breakthrough innovation in food. We believe that an integrated approach of big-data and experiments, can respond to changing consumer demands and sustainability needs with greater agility and efficiency in diverse applications like plant breeding and innovative food formulation.

## Footnotes

3 Note that the fundamental structure, i.e. the branching points and chain lengths is unchanged.

4 “model configuration” refers to a set of predictive features along with a value for each hyperparameter.

5 Though the bin size of 5 was also considered in the low-resolution approach, it was not the optimal bin size for any prediction target.

6 The DP numbers for short and long branches of amylose have been suggested as DP 100 - 700 and 700-40,000 in the proportion of 90% and 10%. In the case of rice, this range has been specified as 270-354 and 1550-1965 by Wang et al^45^.

## Notes

### Competing Interest Statement

The authors have declared no competing interest.

